# Reduced cerebral blood flow and metabolic rate trigger an adaptive mitochondrial remodeling program in a Down syndrome mouse model

**DOI:** 10.64898/2026.10.27.762261

**Authors:** Muzna Saqib, Dylan C. Sarver, Zhiliang Wei, G. William Wong

## Abstract

Mitochondrial dysfunction is a central feature of Down syndrome (DS), yet how trisomy 21 affects brain bioenergetics across scales—from gene transcription to cellular function to whole-brain metabolism— remains poorly understood. Here, we use the Dp16(1)Yey/+ (abbreviated Dp16) DS mouse model, which carries ∼115 triplicated gene orthologs of human chromosome 21, to address this major knowledge gap and to provide direct *in vivo* evidence of brain-wide metabolic deficit in DS. Using magnetic resonance imaging (MRI) approaches, we show that Dp16 mice have reduced cerebral blood flow and metabolic rate of oxygen, with effects more pronounced in males than in females. Despite the suppression of cerebral metabolic rate, mitochondrial respiratory capacity and ATP synthase activity are largely preserved across brain regions, with sex differences noted. Transcriptomic analyses reveal marked upregulation of mitochondrial OXPHOS genes and the convergence of biological pathways related to proteostasis and mitochondrial function across brain regions, with male cortex and female cerebellum showing the biggest change. Collectively, our data suggest that a brain-wide metabolic deficit in Dp16 mice triggers an adaptive mitochondrial remodeling program across the brain to support bioenergetic demands. This compensatory response, however, is insufficient to normalize the metabolic dysfunction in Dp16 mice. Our findings underscore the value of integrating functional assessments at the organismal and tissue levels with transcriptomic changes to gain a systems view of physiologic perturbations and adaptations in the DS brain which may influence phenotypic outcomes.

## INTRODUCTION

Down syndrome (DS), caused by complete or partial trisomy of human chromosome 21 (Hsa21), is the most common genetic cause of intellectual disability, occurring in approximately 1 in ∼800 live births (1). Its neurological consequences extend well beyond cognitive impairment, as virtually all adults with DS develop the neuropathological hallmarks of Alzheimer’s disease by age 40 (2). Additionally, approximately 70% of adults receive a clinical dementia diagnosis by mid-fifties, and lifetime risk approaches 95% (3). DS is increasingly understood as a model of accelerated aging, in which trisomy 21 drives amyloid accumulation, oxidative stress, neuronal loss, and neuroinflammation, establishing a chronically stress-prone cellular environment decades before similar hallmarks appear in age-matched non-DS individuals (4). Central to this phenotype is mitochondrial imbalance, disrupted energy metabolism, elevated oxidative stress, impaired mitochondrial dynamics, and defective quality control mechanisms [reviewed in (5–8)]. However, much of the evidence concerning DS-associated deficits in mitochondrial morphology, dynamics (i.e., fusion/fission), and function are derived from *in vitro* model systems (9–19), with limited studies examining tissue mitochondrial health *in vivo* (20–26).

Despite variability in severity, people with DS generally have reduced brain volume, white matter volume, gyral complexity, sulcal depth, and cortical plate growth; some of these deficits are apparent in fetal and neonatal DS brains (27). Developmental brain abnormalities, along with accelerated brain aging in DS suggest an early, developmentally-timed insult that is carried forward and compounded by a slower, cumulative metabolic and functional decline across the lifespan, as shown by a linear decline in [F-18]-flurodeoxyglucose (FDG) uptake many years before the onset of dementia (28). It is likely that trisomy 21 drives an early mitochondrial dysfunction during neurodevelopment that leaves neurons chronically primed; the cumulative bioenergetic and oxidative burden of aging eventually results in a major decline in brain metabolism and function. This is consistent with the idea that disrupted mitochondrial bioenergetics, redox homeostasis, and organellar quality control in DS contribute to neurodegeneration (4, 5, 29). However, despite recent efforts including positron emission tomography (PET) studies on human brain glucose (FDG) uptake (28, 30, 31), the molecular mechanisms by which trisomy 21 disrupts metabolism and mitochondrial function, particularly in the brain, are still poorly understood.

Given the intrinsic limitations of human studies, we employed the Dp(16)1Yey/+ (abbreviated Dp16) DS mouse model (32) in the present study to determine the impact of triplicated genes on brain metabolism across scales, i.e., from assessing whole-brain metabolism in intact animals to mitochondrial function and transcriptomic changes in different brain regions. The Dp16 mouse model is widely used to study DS-associated phenotypes (32, 33). This model was created using a precise Cre-loxP based chromosomal engineering approach and carries a duplicated segment of mouse chromosome 16 (Mmu16) that is orthologous to Hsa21, with ∼115 triplicated genes (32). In the context of brain function, the Dp16 mouse model recapitulates the neurological and cognitive deficits seen in DS (34–39), thus making it a useful model to investigate the metabolic underpinnings of DS neuropathology.

Here we use a non-invasive MRI method to directly assess whether the Dp16 mice have a brain-wide deficit in metabolism, and to correlate the brain metabolic phenotype to mitochondrial activity and transcriptomics changes across brain regions. Although mitochondrial dysfunction is well documented in cell and animal model systems, whether this translates to a brain-wide deficit in metabolic rate in the intact brain of DS or animal models has not been directly demonstrated. It is worth nothing that PET imaging of people with DS measures only the uptake of FDG—a glucose analog that is trapped in cells after phosphorylation—as a proxy for brain metabolism without directly assessing glucose metabolism (i.e., glycolytic flux) and mitochondrial function (i.e., the use of oxygen to fully oxidize substrates through TCA cycle and OXPHOS). Our multimodal approach allows us to relate cerebral blood flow and metabolic rate of oxygen to mitochondrial bioenergetics and gene expression programs across multiple brain regions. The combination of *in vivo* neuroimaging, *ex vivo* bioenergetic profiling, and transcriptomic analysis have yielded unexpected findings. We showed that the Dp16 mice have reduced cerebral blood flow and metabolic rate of oxygen, consistent with brain hypometabolism as shown by FDG uptake in people with DS (28, 30, 31). However, contrary to expectation, the maximal mitochondrial respiratory capacity across four major brain regions (cortex, hippocampus, cerebellum, and hypothalamus) is largely preserved, along with complex V (ATP synthase) activity, despite some sex differences noted. Our transcriptomic data reveal marked upregulation of genes and pathways related to mitochondrial function and proteostasis. These collective data suggest an adaptive response, at the cellular and transcriptomic levels across brain regions, to metabolic deficits in the Dp16 mice. Despite its mitigating effect, the compensatory mitochondrial remodeling program is not sufficient to normalize the reduced metabolic rate of oxygen in male Dp16 mouse brain. Nevertheless, our findings offer a plausible explanation—depending on the extent of compensation—for the variable severity in phenotypic outcomes in DS (40–42), as well as the differential survival of trisomy fetuses during development (43).

## RESULTS

### Workflow and analysis of brain bioenergetics in the Dp16 mouse model

We used both male and female mice in our study to determine if there are potential sex differences in brain bioenergetics. A sample size of 12 per genotype per sex was used to ensure the study is sufficiently powered and robust (**Fig. 1A**). Our workflow began with *in vivo* MRI scans, which allowed us to acquire data on respiration rate, brain volume, cerebral blood flow (CBF), oxygen extraction fraction (OEF), and cerebral metabolic rate of oxygen (CMRO_2_) (**Figure 1B**). Respiratory rate was recorded during MRI scanning as a physiological indicator in mice. CBF indicates the amount of blood flow supplied to the brain via the feeding arteries. OEF indicates the percentage of oxygen extracted by the brain tissue from capillary bed. CMRO_2_ indicates the consumption rate of oxygen by the brain. Respiration rate was used as a marker of anesthetic depth. In our previous study (44), we found a calibratable relationship between respiration rate and vascular physiology. For this reason, we included respiration rate as a covariate when interpreting CBF, OEF, and CMRO_2_. All mice were at 16 weeks of age when MRI scans were performed. Following MRI scans, mice underwent post-imaging dissections for brain tissue collection (**Figure 1C**). The four brain regions (cortex, hippocampus, cerebellum, and hypothalamus) were used for two parallel downstream analyses. First, the maximal mitochondrial respiratory capacity was assessed using a Seahorse-based platform to determine oxygen consumption rate (OCR) and acidification rate (AR) linked to individual electron transport chain complexes, as well as ATP synthase activity (**Figure 1D**). Second, the same brain regions were subjected to bulk RNA sequencing to evaluate changes in transcriptome and biological pathways impacting brain bioenergetics and mitochondrial function (**Figure 1E**). Together, our multimodal approaches aimed to relate *in vivo* brain metabolism in an intact animal to changes in mitochondrial activity and transcriptome across four major brain regions in a DS mouse model.

**Figure 1.**
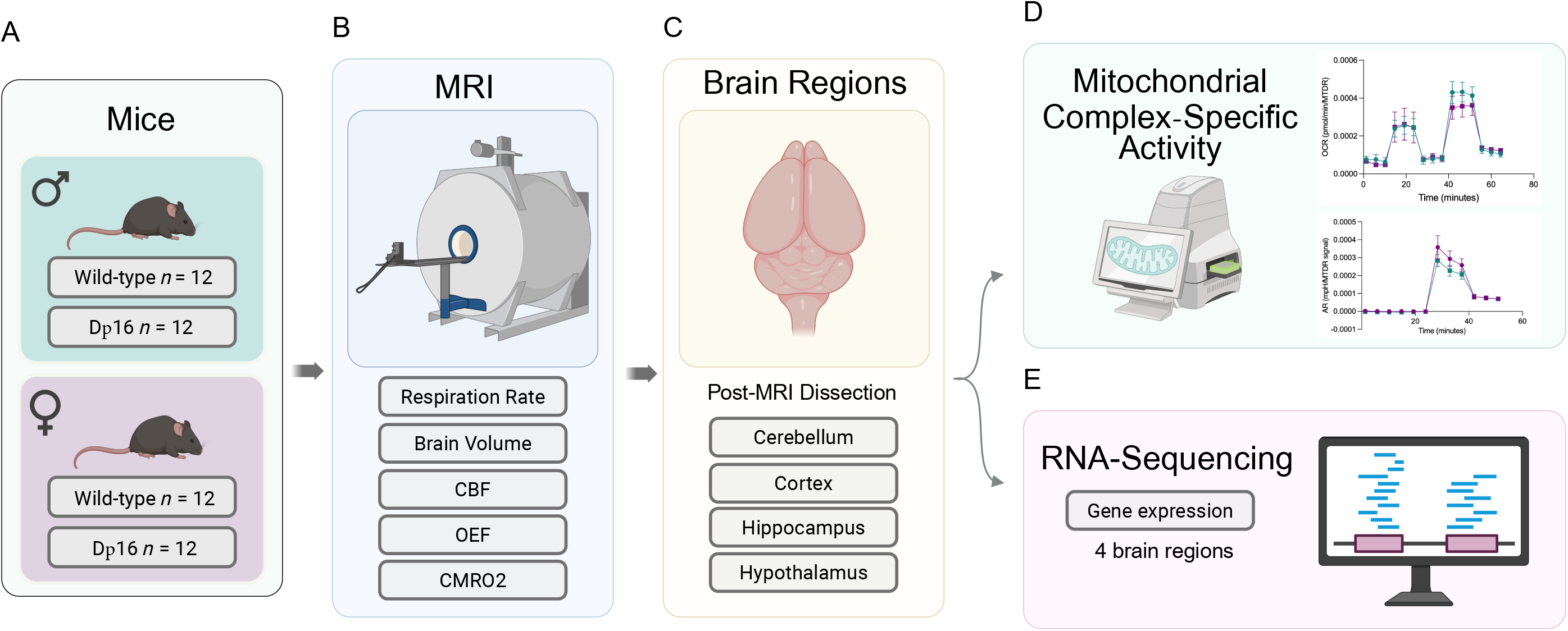
Workflow and analysis of brain bioenergetics in the Dp16 mouse model. (A) Diagram showing the sample sizes of male and female mice for WT and Dp16 groups. (B) Schematic of MRI acquisition and the parameters collected. CBF, cerebral blood flow; OEF, oxygen extraction fraction; CMRO2, cerebral metabolic rate of oxygen. (C) Brain regions collected following dissection, including the cerebellum, cortex, hippocampus, and hypothalamus. (D) Isolated brain regions were used to assess mitochondrial complex I to V activity via high-resolution mitochondrial respirometry. (E) Transcriptomic analyses of the same brain regions were carried out to evaluate changes in gene expression and biological pathways impacting brain bioenergetics.

### Reduced cerebral blood flow and metabolic rate of oxygen in Dp16 mice

We used T2-relaxation-under-spin-tagging (TRUST) and phase contrast (PC) MRI scans to obtain quantitative information on cerebral venous oxygenation and blood flow, respectively (details are provided in the methods section). TRUST MRI generated control and labeled images centered on the sagittal sinus, with the corresponding difference image used to quantify blood T2 and, in turn, oxygen extraction fraction (OEF) (**Figure 2A**). Representative difference images showed a clear signal at the sinus confluence across WT and Dp16 male and female mice. In parallel, PC MRI was used to generate vector maps of the left and right internal carotid arteries (LICA and RICA) and the basilar artery (BA), from which blood flow velocity was calculated (**Figure 2B**), with vessel signal clearly distinguishable across all three vascular territories in WT and Dp16 male and female mice.

**Figure 2.**
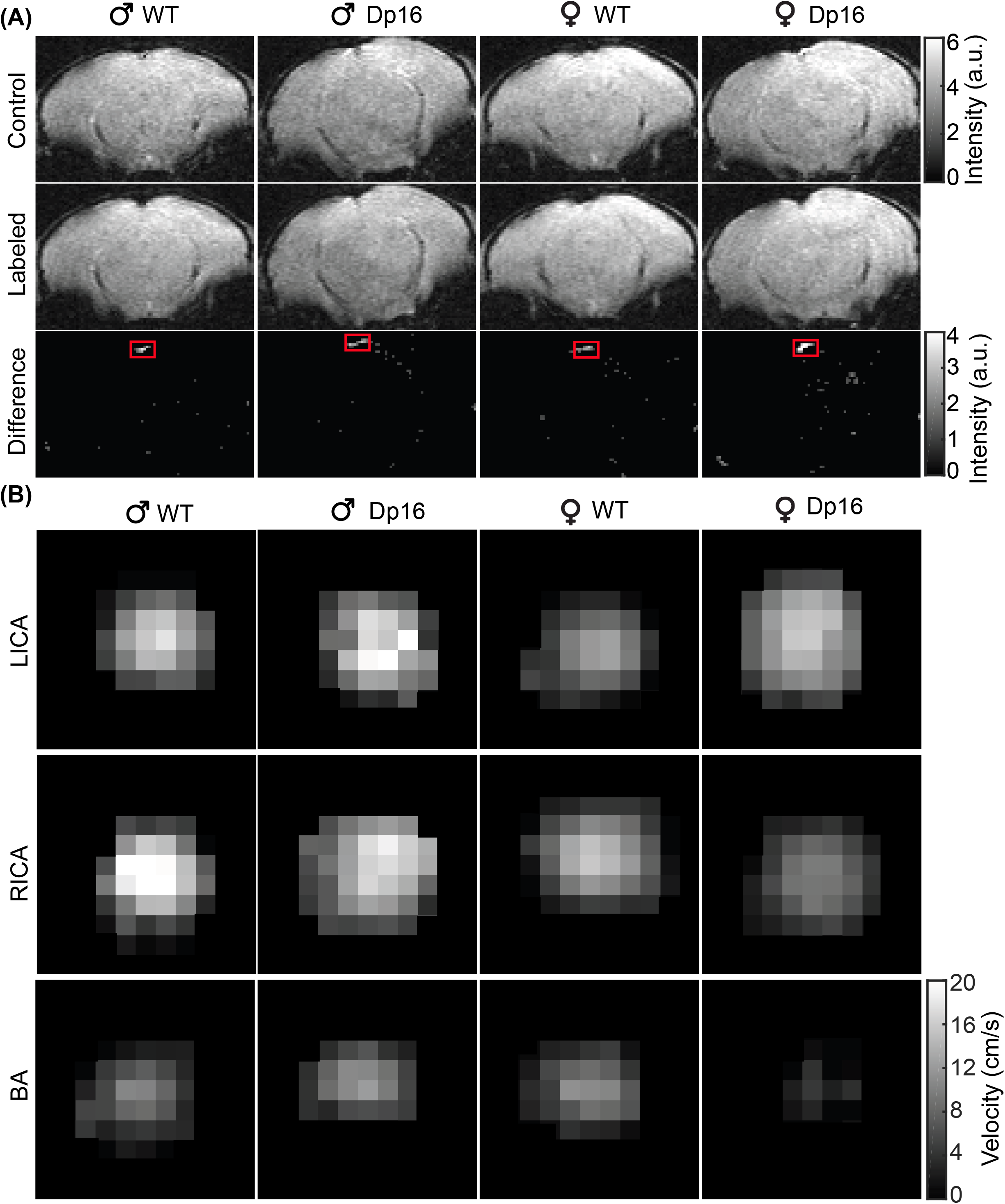
Representative results of T2-Relaxation-Under-Spin-Tagging (TRUST) and phase contrast (PC) magnetic resonance imaging (MRI). A) Representative control, labeled, and difference images at the effective echo time of 0.13 ms from TRUST MRI, showing the sinus confluence (red squares) in male and female WT and Dp16 mice at 16 weeks of age. (B) Velocity maps of the left internal carotid arteries (LICA, upper panel), right internal carotid arteries (RICA, middle panel), and basilar artery (BA, lower panel). Display range of blood-flow velocity is 0-20 cm/s. For each mouse, blood flows through the LICA, RICA, and BA were summed to estimate CBF.

Using this approach, we compared respiration rate, brain volume, cerebral blood flow (CBF), oxygen extraction fraction (OEF), and cerebral metabolic rate of oxygen (CMRO_2_) between WT and Dp16 mice (**Figure 3**). In Dp16 males, CBF and CMRO_2_ were significantly reduced relative to WT controls, while respiration rate, brain volume, and OEF were not different from WT control (**Figure 3A-E**). In Dp16 females, brain volume was modestly larger relative to WT controls (**Figure 3 G**). While respiration rate and OEF were not different between genotypes, CBF and CMRO_2_ were lower in Dp16 females, though not significantly, when compared to WT controls (**Figure 3F, H-J**). Together, these data show for the first time that cerebral blood flow and metabolic rate of oxygen are reduced in a DS mouse model.

**Figure 3.**
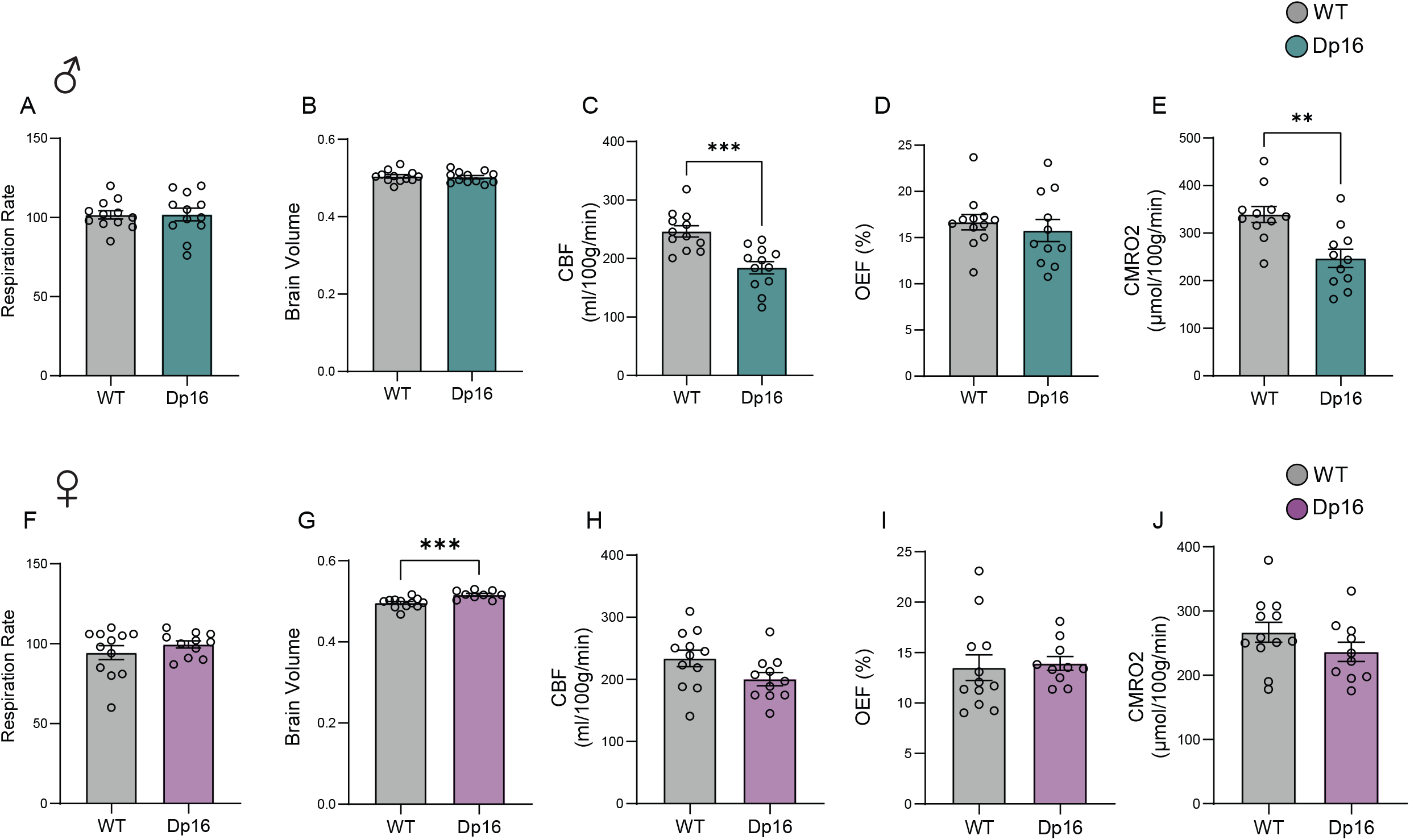
Reduced cerebral blood flow and metabolic rate in Dp16 mice. (A, F) Respiration rate, (B, G) brain volume, (C, H) cerebral blood flow (CBF), (D, I) oxygen extraction fraction (OEF), and (E, J) cerebral metabolic rate of oxygen (CMRO_2_) in wild-type (WT) and Dp16 males (A–E) and females (F–J). Data are presented as mean ± SEM with individual data points shown; *n* = 12 mice/sex/genotype. **p < 0.01, ***p < 0.001 by Welch’s *t*-test

### Mitochondrial respiratory capacity across the brain is largely preserved in Dp16 mice

Most of the oxygen consumed (∼90%) is used by the mitochondria to oxidize glucose, fatty acids, and amino acids through the TCA cycle and OXPHOS for ATP production (45). We next asked whether the reduced metabolic rate of oxygen in Dp16 mice is due to impaired mitochondrial function. We used a high-resolution mitochondrial respirometry approach to interrogate mitochondrial complex I (CI), CII, CIV, and CV (ATP synthase) activity in frozen tissue samples (46, 47). Different electron donors were used to assess the activity of different complexes: NADH for CI, succinate for CII, TMPD and ascorbate for CIV. Measurements of oxygen consumption rate (OCR) through CI, CII or CIV provide quantitative information on mitochondrial function and health. All OCR data were normalized to mitochondrial content, as quantified by MitoTracker Deep Red (MTDR) signals (**Figure 4 – figure supplement 1**). We did observe reduced mitochondrial content in Dp16 male hypothalamus and increased mitochondrial content in Dp16 female cortex and hypothalamus. To assess CV (ATP synthase) hydrolytic activity, ATP was provided and the acidification rate (AR) was measured. We showed that Dp16 males had higher OCR through CI in the cerebellum, lower OCR at CIV in the hippocampus, and higher ATP synthase hydrolytic activity in the cortex, hippocampus, and cerebellum (**Figure 4**). In Dp16 females, OCR through CI, CII, and CIV were not different from WT controls, whereas ATP synthase hydrolytic activity was lower in the cortex and cerebellum (**Figure 4**). Together, these data indicate that while there are some sex differences, particularly related to ATP synthase activity, most of the parameters were not different between genotypes of either sex, indicating that the maximal mitochondrial respiratory capacity across the brain is largely preserved in Dp16 mice (21 weeks old) despite reduced cerebral blood flow and metabolic rate of oxygen.

**Figure 4.**
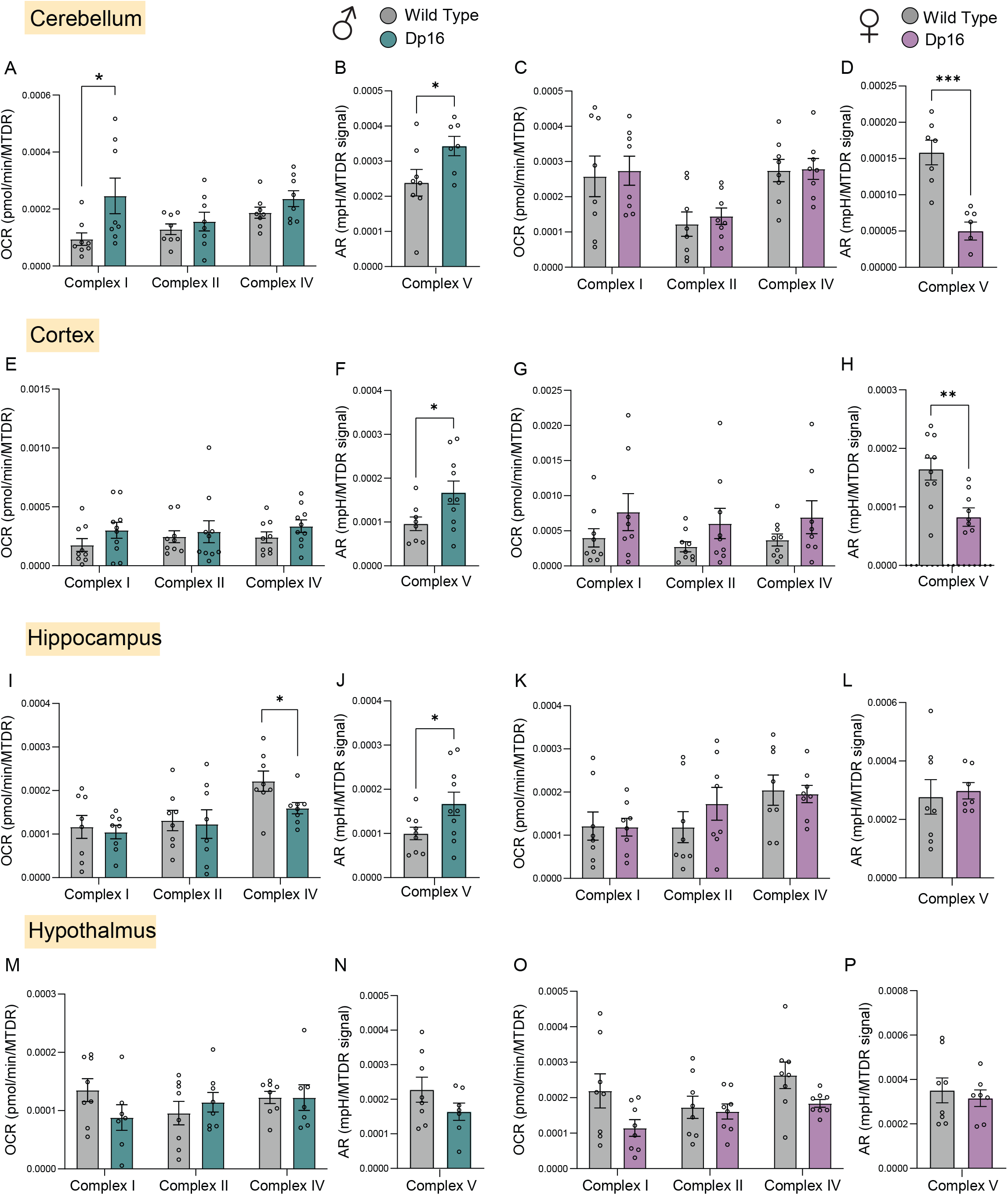
Mitochondrial respiratory capacity is largely preserved across brain regions in Dp16 mice. Mitochondrial respiration through complex I (CI, NADH-stimulated), complex II (CII, succinate-stimulated in the presence of rotenone, a CI inhibitor), and complex IV (CIV, via TMPD + ascorbate), and ATP hydrolysis activity (CV, ATP-stimulated), in male cerebellum (A,B), female cerebellum. (C,D), male cortex (E,F), female cortex (G,H), male hippocampus (I,J), female hippocampus (K,L), male hypothalamus (M,N), and female hypothalamus (O,P). All oxygen consumption rates (OCR) and acidification rates (AR; a measure of ATP hydrolysis rate) are normalized to mitochondrial content (based on MTDR). Data are presented as mean ± SEM with individual data points shown; each biological replicate represents the average of three technical replicates. Sample size per group: WT = 8 and Dp16 = 8. **p < 0.01, ***p < 0.001 by Welch’s *t*-test.

### Increased expression dosage of triplicated genes across the brain in Dp16 mice

We next performed RNA sequencing across the same brain regions to gain molecular insights into the brain metabolism of Dp16 mice. The Dp16 model carries ∼115 triplicated genes (32). We previously showed that the majority of triplicated genes are expressed at 1.5-fold or higher in peripheral tissues (e.g., liver, fat, muscle), in accordance with increased gene dosage (21). Similarly, in the brain, the majority of triplicated genes in cortex, hippocampus, cerebellum, and hypothalamus were also expressed at ∼1.5-fold or higher (**Figure 5A-B**). In female but not male cerebellum and hippocampus, however, the expression of multiple triplicated genes were suppressed relative to WT controls, thus highlighting tissue- and sex-specific variations in the expression patterns of triplicated genes; such variegated expression patterns of chromosome 21 genes have been previously documented in people with DS (48). Similar to what we have seen in the peripheral tissues (21), there was sex-biased expressions for some of the triplicated genes that may influence phenotypic outcomes between the sexes (**Figure 5C**). Together, these data provide a gene expression atlas for the triplicated genes across four major brain regions in Dp16 male and female mice, with notable sex differences.

**Figure 5.**
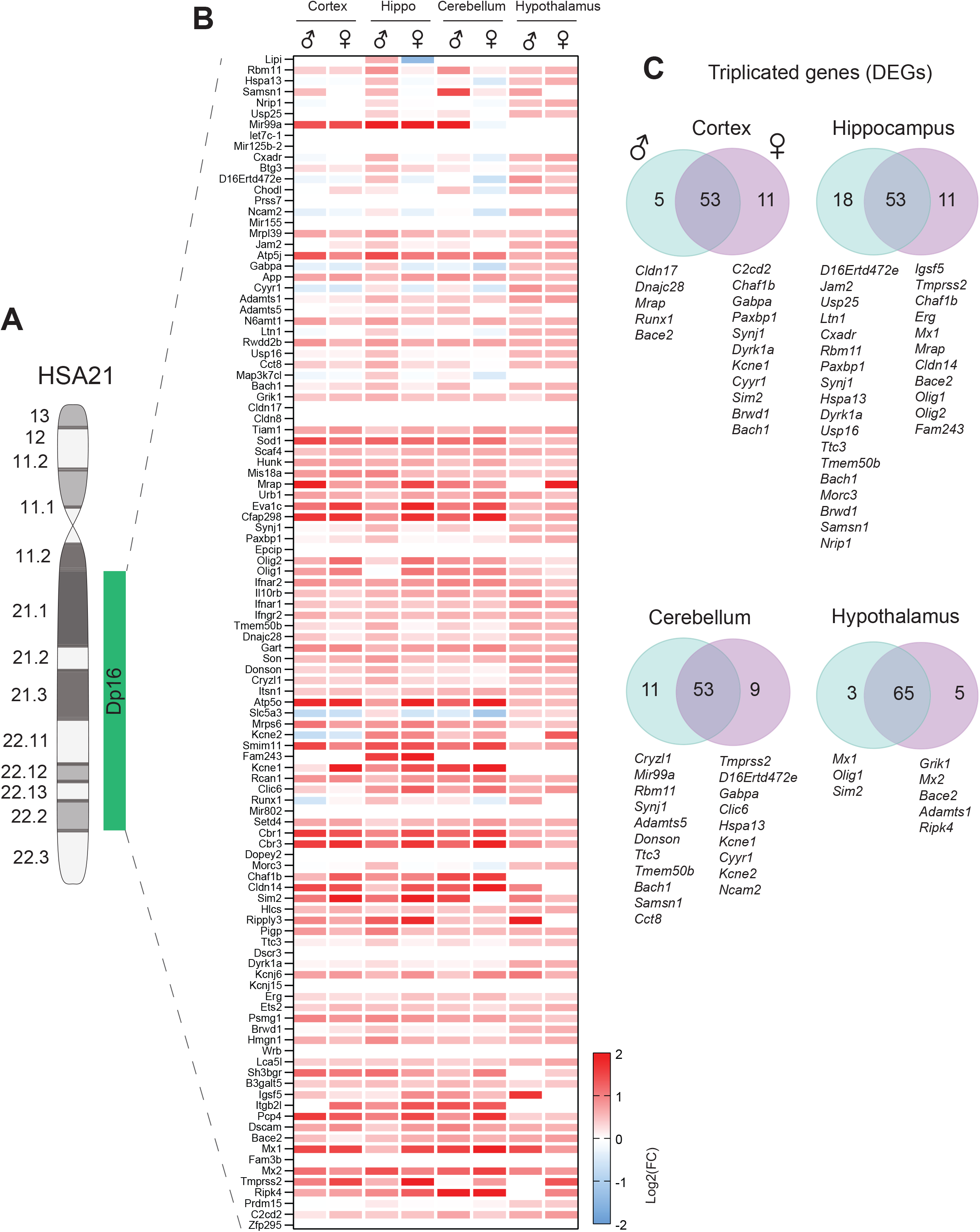
Increased expression dosage of triplicated genes across brain regions in Dp16 male and female mice. (A) Graphical representation of human chromosome 21 (Hsa21) and the syntenic Mmu16 segment that is triplicated in Dp16 mice. (B) Heatmap of gene expression changes (log2 fold change, Dp16 vs. WT) for genes within the triplicated interval, shown separately for male and female mice across cortex, hippocampus, cerebellum, and hypothalamus. Red indicates increased expression and blue indicates decreased expression relative to WT (C) Venn diagrams showing the overlap between male and female differentially expressed genes (DEGs) among triplicated genes within each brain region, with sex-specific DEGs listed below each diagram. DEG is defined as log2(FC) > 0 with an adjusted p-value (Padj) < 0.05. Sample size. *n* = 8 per genotype per sex per tissue-type (cortex, cerebellum, hippocampus). For hypothalamus, *n* = 6 per genotype per sex.

### The triplicated genes alter the brain transcriptome in a sex- and region-specific manner

Next, we determined the impact of triplicated genes on the transcriptome across brain regions when mice were 18 weeks of age. Across all brain regions, there were more upregulated differentially expressed genes (DEGs) than downregulated (**Figure 6A**). In Dp16 males, the cortex has the largest number of DEGs, followed by cerebellum, hippocampus, and hypothalamus. In Dp16 females, cerebellum has the largest number of DEGs, followed by hippocampus, cortex, and hypothalamus. Surprisingly, only modest overlap was seen between DEGs across sex, with the majority of the upregulated or downregulated DEGs showing sex-biased expression across brain regions (**Figure 6B**), a finding that also holds true in peripheral tissues (21). We performed pathway enrichment analyses using Gene Ontology (49) to see what biological processes are dysregulated in the brain. To our surprise and contrary to expectation, many of the top enriched pathways upregulated in male cortex, male and female hippocampus, and female cerebellum are related to mitochondrial function, cellular respiration, and oxidative phosphorylation (**Figure 6C**). In contrast, the top upregulated pathways in hypothalamus are related to immune and cytokine signaling in Dp16 males and extracellular matrix remodeling in Dp16 females. The top downregulated pathways in cerebellum are related to ion channels (e.g., GABA-gated chloride channel), transporter and receptor activity in females, and GABAergic synaptic transmission and postsynaptic membrane in males. In the hypothalamus, the top downregulated pathways are related to synaptic signaling and transmission in females, and mRNA processing and translation in males. Together, these data highlighted major sex differences in transcriptome across brain regions, along with shared and distinct biological pathways impacted by the triplicated genes.

**Figure 6.**
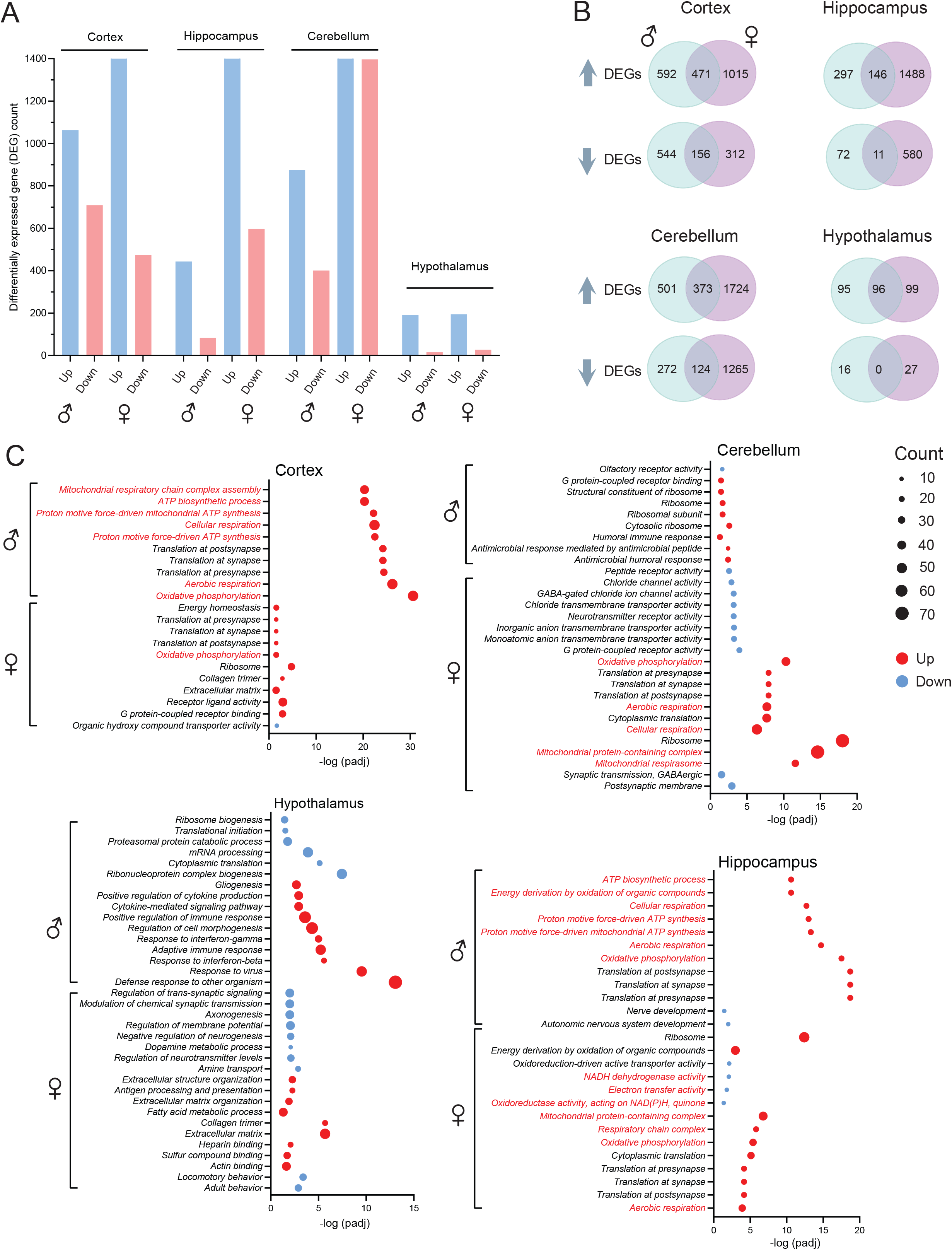
Altered transcriptome and biological pathways in Dp16 cortex, hippocampus, cerebellum, and hypothalamus. (A) Number of differentially expressed genes (DEGs) up- and down-regulated in males and females across four brain regions. For the cortex, hippocampus, and cerebellum, DEG is defined as log2(FC) > 1 and FDR adjusted p-value (Padj) < 0.05. For the hypothalamus, DEG is defined as log2(FC) > 0.5 and Padj < 0.05. (B) Venn diagrams showing the overlap between male and female DEGs, plotted separately for upregulated and downregulated genes in each brain region. (C) Gene Ontology (GO) enrichment analysis of biological pathways altered in each brain region, shown separately for males (top) and females (bottom). The top enriched GO terms are plotted by significance [-log(padj)], with dot size indicating gene count and color indicating direction of change (red = upregulated, blue = downregulated).

### Upregulation of mitochondrial OXPHOS genes across the brain in Dp16 mice

Given the shared upregulated pathways related to mitochondrial function across brain regions, we decided to examine more closely all the genes that encode mitochondrial OXPHOS complexes (CI to CV). Except the hypothalamus, most of the nuclear-encoded OXPHOS genes are upregulated, whereas the mitochondrial genome-encoded CI component (*mt-nd2*, *mt-nd3*, *mt-nd4*, *mt-nd4l*, *mt-nd5*, *mt-nd6*) are suppressed, in the cortex, hippocampus, and cerebellum of Dp16 mice, with the exception of the male hippocampus, in which several of the genes were upregulated (**Figure 7A**). While most of the changes in OXPHOS gene expression were shared between Dp16 males and females, there were several OXPHOS genes that showed region-specific discordant expression patterns between the sexes; for instance, a subset of genes showed an opposite directionality of change in the cortex, hippocampus, and cerebellum, whereas another subset of genes showed an asymmetric change in the hypothalamus (i.e., up-or down-regulated in one sex with no change in the other sex) (**Figure 7B**). Since we observed reduced cerebral metabolic rate of oxygen, the coordinated upregulation of most mitochondrial OXPHOS genes suggests an adaptive response across brain regions to mitigate impaired whole-brain metabolism in the Dp16 mice.

**Figure 7.**
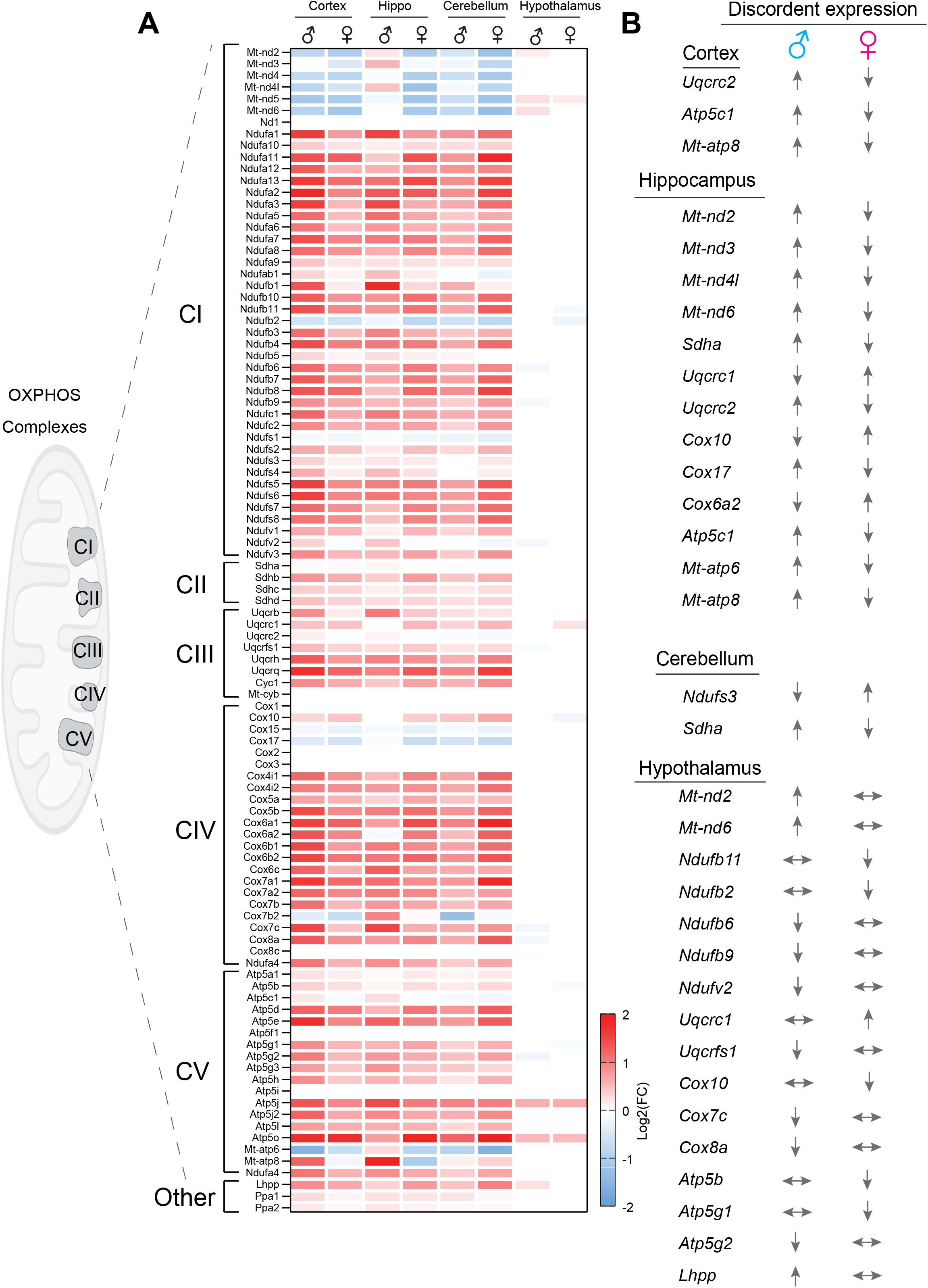
Majority of OXPHOS genes are upregulated across brain regions in Dp16 mice. (A) Heatmap of gene expression changes (log_2_(fold change), Dp16 vs. wildtype) for genes encoding subunits of mitochondrial complex I–V and related OXPHOS machinery, organized by complex (CI–CV and other) and shown separately for male and female mice across cortex, hippocampus, cerebellum, and hypothalamus. Red indicates increased expression and blue indicates decreased expression relative to WT controls. (B) Genes showing discordant (opposite-direction) expression changes between males and females within each brain region, with arrows indicating the direction of change in each sex; double-headed horizontal arrows indicate no significant change.

### Mitochondrial remodeling across the brain in Dp16 mice

To provide further evidence of mitochondrial adaptations at the transcriptomic level, we conducted a mitochondrial module analysis to identify regional and sex-specific patterns of mitochondrial-associated transcriptional remodeling in Dp16 mice. We focused on nine modules related to mitochondrial function; these include the respiratory chain and associated mitochondrial pathways, including electron transport chain complexes (CI, CII, CIII, CIV, CV), coenzyme Q (Q-linked), cytochrome c (Cyt c), the tricarboxylic acid cycle (TCA), and assembly factors (AF). Module scores were calculated for each individual mouse by averaging the normalized expression values of all genes assigned to the corresponding module. The change in module (**Δ**module) score represents the magnitude of change between Dp16 and WT mice for a given module. Our analyses showed that the Dp16 male cortex and female cerebellum represent the brain regions with the greatest difference compared to WT controls (**Figure 8A and Figure 8 – figure supplement 1**). This indicates that sex influences which brain regions undergo the greatest level of mitochondrial-associated gene expression remodeling.

**Figure 8.**
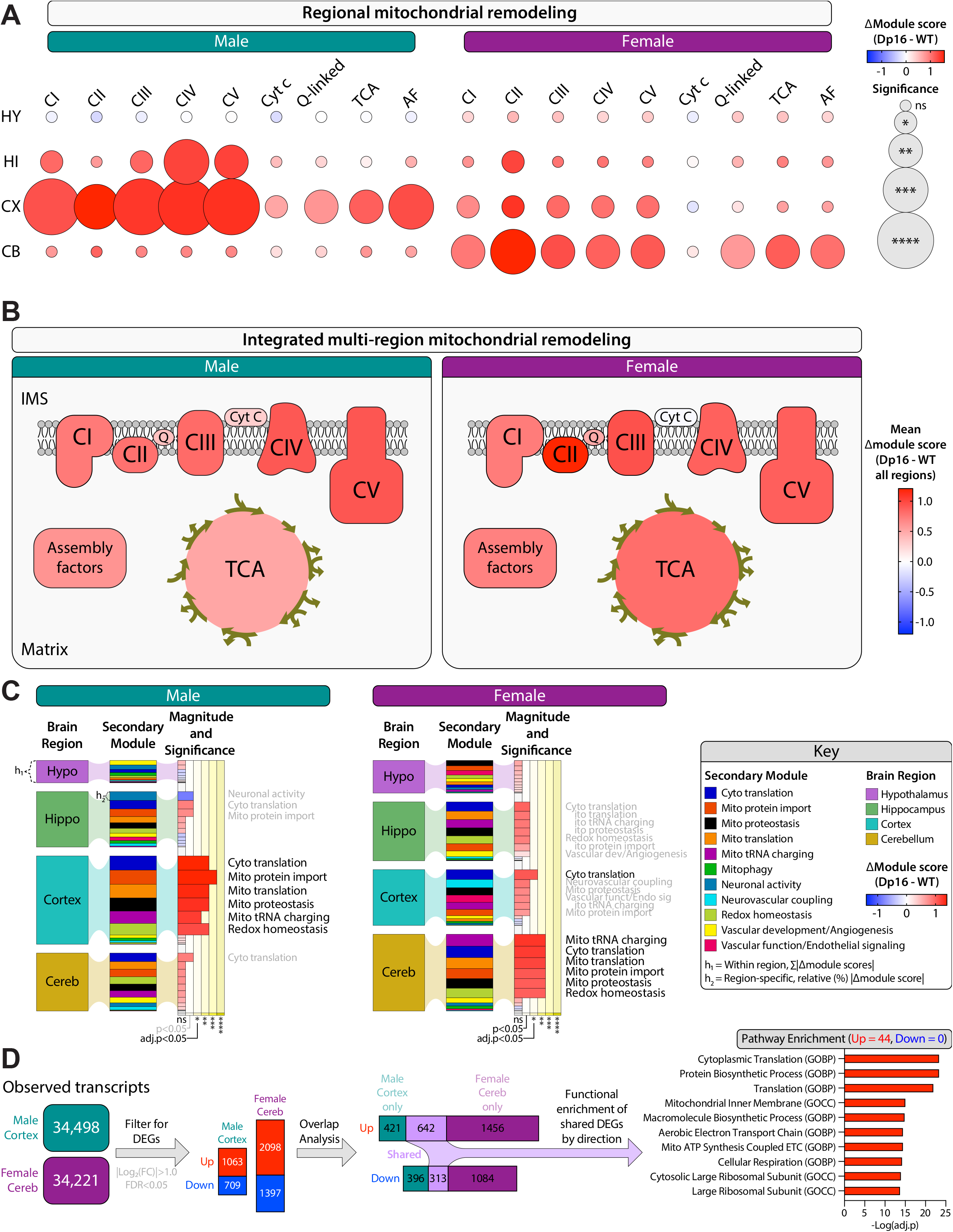
Sex- and region-specific mitochondrial transcriptional remodeling is accompanied by coordinated and conserved transcriptional programs in the Dp16 brain. (A) Bubble plot of mitochondrial module analysis across hypothalamus (HY), hippocampus (HI), cortex (CX), cerebellum (CB), and sex, showing the change in module score (Δmodule score, Dp16 vs. WT) for electron transport chain complexes (CI–CV), coenzyme Q (Q-linked), cytochrome c (Cyt c), the TCA cycle, and assembly factors (AF). Red indicates Dp16 > WT and blue indicates Dp16 < WT; circle size denotes statistical significance. (B) Integrated multi-region analysis showing the mean Δmodule score (averaged across all brain regions) for each mitochondrial module by sex. (C) Secondary module analysis showing magnitude and significance of Δmodule score for functional programs related to mitochondrial and cellular maintenance by brain region and sex. (D) Expression overlap and functional enrichment analysis of genome-wide differentially expressed genes (DEGs, |Log2FC|>1.0, adj. p<0.05) shared between male cortex and female cerebellum, the two regions with the greatest mitochondrial remodeling, separated by direction of change, with pathway enrichment shown for shared upregulated DEGs. Abbreviation: Hypo, hypothalamus; Hippo, hippocampus; Cereb, cerebellum.

To provide a composite view of mitochondrial module changes in Dp16 mice, we performed integrated multi-region analyses of mitochondrial remodeling across the brain. The mean module was calculated by averaging each mitochondrial module across all brain regions within a sex. The mean **Δ**module score represents the magnitude of change between Dp16 and WT mice for a given integrated module expression level. These analyses showed that regardless of sex, mitochondrial-associated gene expression is elevated across the Dp16 brain in nearly all nine modules analyzed (**Figure 8B**).

We next performed a secondary module analysis to further contextualize the mitochondrial remodeling by examining additional biological processes that may contribute to or accompany changes in mitochondrial function. These eleven secondary modules include cytoplasmic translation, mitochondrial protein import, mitochondrial proteostasis, mitochondrial translation, mitochondrial tRNA charging, mitophagy, redox homeostasis, neuronal activity, neurovascular coupling, vascular development/angiogenesis, and vascular function/endothelial signaling. Our analyses showed that elevated mitochondrial module expression levels are accompanied by increased secondary module expression for cytosolic translation, mitochondrial protein import, mitochondrial translation, mitochondrial proteostasis, mitochondrial tRNA charging, and redox homeostasis, in both male and female mice (**Figure 8C and Figure 8 – figure supplement 2-3**). This suggests that mitochondrial-associated transcriptional remodeling in the Dp16 brain is accompanied by a conserved program of mitochondrial maintenance and cellular biosynthetic processes.

### Conserved mitochondrial-associated transcriptional programs in the Dp16 brain

Lastly, we performed genome-wide differential expression analysis to compare male cortex and female cerebellum to identify transcriptional features shared between these two brain regions with the highest level of mitochondrial-associated transcritional remodeling (**Figure 8D**). From the 642 shared upregulated and 313 shared downregulated genes between male cortex and female cerebellum, 44 significantly enriched pathways were identified among the shared upregulated genes, whereas no pathways reached significance among the shared downregulated genes after multiple testing correction. Our pathway enrichment showed that Dp16 brain regions with elevated mitochondrial remodeling share genes involved in translation, protein biosynthesis, and mitochondrial energy production pathways regardless of sex. Because the preceding analyses were focused on predefined mitochondrial and related functional modules, we next examined the brain transcriptome without restricting our analysis to mitochondrial genes or pathways. This independent, genome-wide analysis revealed substantial overlap in the transcriptional programs of the male cortex and female cerebellum, despite their sex- and region-specific magnitudes of mitochondrial remodeling. There was little overlap between genes with opposing directionality of change (i.e., male cortex down vs female cerebellum up). Together, these data suggest that the Dp16 brain engages a conserved set of mitochondrial-associated transcriptional programs.

Collectively, these analyses highlighted sex- and region-specific mitochondrial remodeling alongside a broader transcriptional program that is conserved across affected brain regions in the Dp16 mice. Across sex and brain regions, mitochondrial function-associated gene expression is consistently elevated, accompanied by increased expression of genes involved in mitochondrial maintenance, protein synthesis, and related biosynthetic processes. These findings argue that the Dp16 brain adapts to an altered energetic state through coordinated up-regulation of mitochondrial and protein synthetic machinery, suggestive of a compensatory response to reduced energetic efficiency and/or increased energetic demands.

## DISCUSSION

Here we use a multimodal approach to provide an integrated view of brain bioenergetics in the Dp16 mice (summarized in **Figure 9**). Our study provides the first direct demonstration of reduced cerebral blood flow and metabolic rate of oxygen in an animal model of DS. Our *in vivo* data is consistent with known clinical findings, in which people with DS have reduced cerebral blood flow (50–55) and brain FDG uptake as a proxy for glucose metabolism (28, 30, 31). Given the intrinsic limitations of performing mechanistic and molecular studies on human subjects, our findings that the Dp16 mice recapitulate brain hypoperfusion and hypometabolism in DS suggest that this is a valuable mouse model to dissect the molecular underpinnings of cerebrovascular and metabolic deficits in DS.

**Figure 9.**
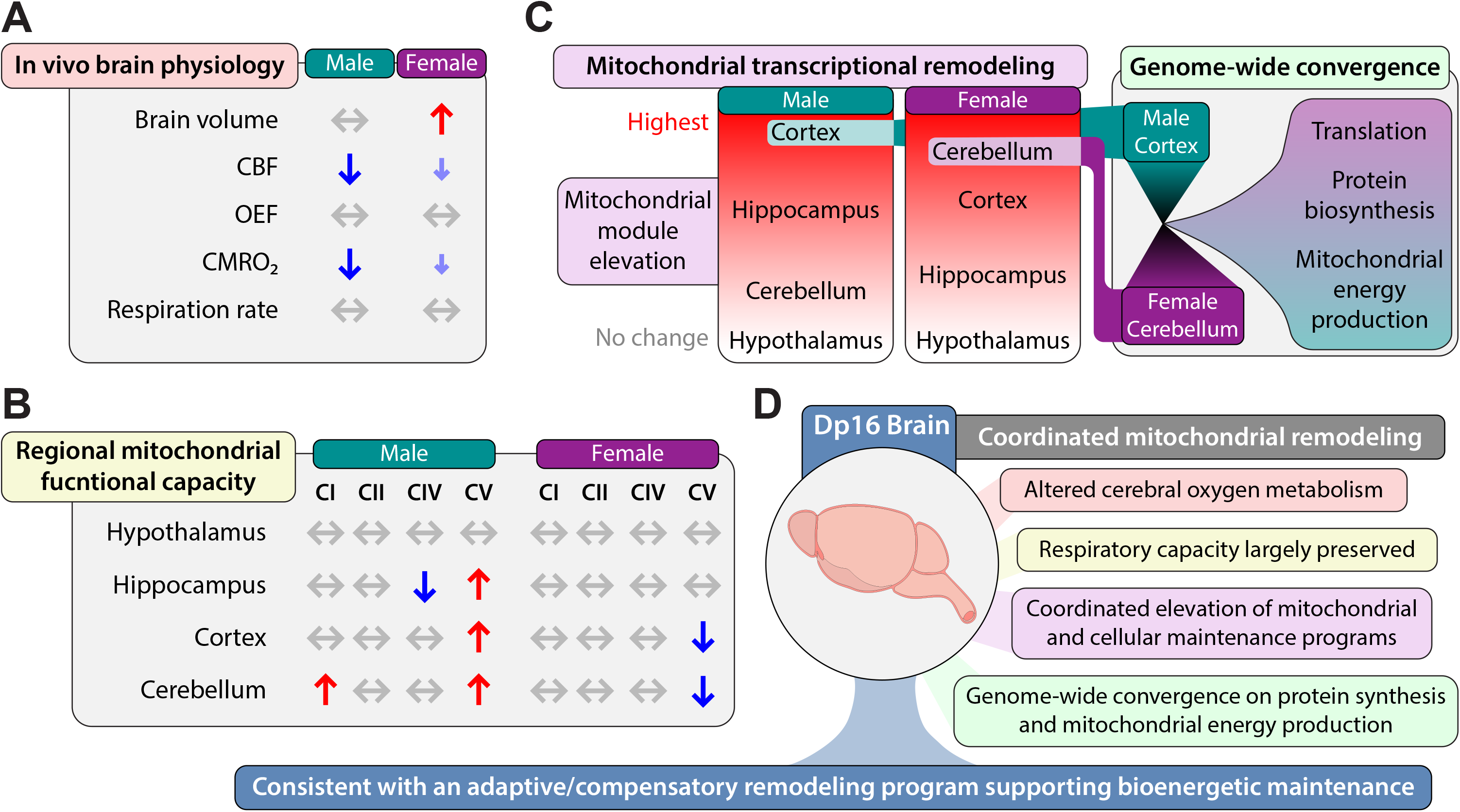
Integrated summary of brain bioenergetic and mitochondrial remodeling in Dp16 mice. Schematic integration of *in vivo* brain physiology (A), regional mitochondrial respiratory capacity (B), and transcriptomic remodeling in the Dp16 brain (C). (A) MRI measurements demonstrate reduced cerebral blood flow (CBF) and metabolic rate of oxygen (CMRO_2_) in male Dp16 mice, with female mice showing similar directional changes that do not reach statistical significance; oxygen extraction fraction (OEF) was unchanged in both sexes. (B) Maximal mitochondrial respiratory capacity is largely preserved across brain regions, with selective sex- and regional-specific alterations in individual respiratory complexes. (C) Transcriptomic analyses identify broad mitochondrial remodeling across the brain, including coordinated changes in respiratory chain, coenzyme Q, cytochrome c, and TCA-associated mitochondrial programs, accompanied by alterations in mitochondrial translation, protein import, proteostasis, tRNA charging, and redox homeostasis. Independent genome-wide analysis of the male cortex and female cerebellum further identified a shared transcriptional program enriched for protein synthesis and mitochondrial energy production. (D) Although the magnitude and regional localization of these changes differ by sex, the integrated findings support a model in which the Dp16 brain undergoes coordinated mitochondrial remodeling in the setting of altered cerebral energetics. This remodeling is consistent with an adaptive or compensatory program that may contribute to mitigating the brain bioenergetic deficits induced by gene dosage imbalance.

To integrate the physiological, functional, and transcriptomic findings, we compared *in vivo* cerebral oxygen consumption with regional mitochondrial respiratory capacity and mitochondrial-related gene expression in the Dp16 brain. Our *in vivo* MRI revealed reduced cerebral blood flow and metabolic rate of oxygen in male Dp16 mice, whereas females showed similar directional changes that did not reach statistical significance (**Figure 9A**). Cerebral blood flow is tightly coupled to metabolic demand (56), a phenomenon known as neurovascular coupling (57) that also underlies the blood-oxygenation-level-dependent (BOLD) effect (58). When brain metabolic rate decreases, CBF is reduced proportionally to match the lower metabolic demand. Since Dp16 males have normal OEF but reduced CMRO_2_, this indicates that the primary driver of reduced CBF is the reduction in CMOR_2_.

Despite a reduction in brain oxygen consumption, maximal mitochondrial respiratory capacity measured *ex vivo* was largely preserved across brain regions (**Figure 9B**). Individual components (i.e., CI to CV) of the respiratory system showed selective sex- and region-specific changes, but there was no evidence of a generalized loss of mitochondrial respiratory capacity. In parallel, transcriptomic analysis revealed broad remodeling of mitochondrial gene expression, with predominantly elevated mitochondrial pathway module scores across brain regions and sexes. This remodeling extended beyond core mitochondrial pathways to include mitochondrial translation, protein import, proteostasis, tRNA charging, and redox homeostasis (**Figure 9C**).

To determine whether these patterns extended beyond predefined modules related to mitochondrial function, we independently examined genome-wide differential gene expression in the male cortex and female cerebellum, the two brain regions exhibiting the greatest mitochondrial transcriptional remodeling in their respective sexes. Concordant differential expression identified 642 shared upregulated and 313 shared downregulated genes. The shared upregulated genes were enriched for protein synthesis, translation, and mitochondrial energy production, whereas no significantly enriched terms were identified among the shared downregulated genes. Together, these findings indicate that the Dp16 brain undergoes coordinated mitochondrial and cellular transcriptional remodeling in the context of altered cerebral oxygen metabolism and largely preserved mitochondrial respiratory capacity. The integrated pattern is consistent with an adaptive or compensatory remodeling program that may contribute to the maintenance of brain bioenergetic capacity in the Dp16 mouse brain (**Figure 9D**).

The notion of compensation, as suggested by our data and pointed out by others (59), has implications for understanding the variable expressivity and severity of DS phenotypes (40–42). While trisomy 21 is the most survivable aneuploidy in humans, a relatively high percentage of trisomic fetuses are spontaneously aborted during pregnancy, presumably due to neurodevelopmental deficits incompatible with life (43). It is also well documented that some DS phenotypes are present in all people with DS, albeit with variable severity; these include cognitive deficits, craniofacial dysmorphology, hypotonia, and early-onset Alzheimer’s-type dementia (1). In contrast, other DS phenotypes are found in some, but not all, people with DS (e.g., congenital heart defects, blood cancers, sensory abnormalities, and increased risk of autoimmune disease). The underlying basis for the variable expressivity and severity of DS phenotypes is largely unknown. We would argue that variable magnitude of compensatory response at the molecular, cellular, and physiological levels across tissues likely contributes to differential survival of trisomic fetuses, as well as the variable severity of DS phenotypes in the postnatal period.

Among the triplicated Hsa21 gene orthologs in Dp16 mice, several are known to play a role in mitochondrial function; for example: *Atp5j* and *Atp5o* encode subunits of mitochondrial ATP synthase (60); *Gabpa* encodes a transcription factor that drive the expression of nuclear-encoded OXPHOS genes (61); *Mrpl39* encodes mitochondrial ribosomal subunits required for translating mtDNA-encoded OXPHOS proteins (62); *Mrps6* encodes a structural component of the mitoribosome (63); *Sim2* encodes a protein that directly interacts with the mitochondrial respiratory chain to facilitate functional supercomplex formation (64). However, it is unclear how the increased expression dosage of these genes positively or negatively affects mitochondrial physiology. Functionally, the mitochondrial respiratory capacity across the brain was largely unaffected in the Dp16 mice. The mitochondrial respirometry assay requires supercomplexes of the respiratory chain to be intact in order for electrons donated by the substrates (NADH and succinate), which enter through CI or CII, to be passed from CoQ, cytochrome C, to the final electron acceptor (CIV); these supercomplexes are preserved in previously frozen tissue (65). Since mitochondrial activity through CI, CII, or at CIV was largely not affected in the Dp16 mice, we therefore infer that the architecture of the mitochondrial supercomplexes is likely to be unaffected as well, though we cannot completely rule out subtle changes in the assembly of the supercomplexes.

There are several limitations of the study we wish to highlight. First, all the data were obtained from mice that are considered relatively young (16-21 weeks of age). A longitudinal study of Dp16 mice would yield valuable information concerning the trajectory of brain bioenergetics across the lifespan and allow us to determine whether Dp16 model recapitulates the progressive and linear decline of brain metabolism with age, as has been shown in individuals with DS (28). Secondly, unlike PET scans in humans which can resolve FDG uptake across different brain regions, the TRUST MRI method we used in mice can only yield a single whole-brain average value of CMOR_2_ and currently lacks the resolution to resolve regional heterogeneity in oxygen consumption. In people with DS, FDG uptake is not reduced uniformly across the brain; suppression of FDG uptake is consistently found in posterior cingulate and parietotemporal cortex (28, 30, 31). Thus, it would be valuable to know whether analogous brain regions are similarly affected in the DS mouse models. Thirdly, our mitochondrial respiration assays were performed on previously frozen tissues. While this method is robust, reproducible, and scalable across many samples (47, 65), it only allows us to measure maximal mitochondrial respiratory capacity in non-intact mitochondria. Fourthly, RNA sequencing data were obtained from bulk tissues; as such, we cannot resolve cell-type-specific contributions to changes in gene expression, even though both neurons and glia are known to be affected in DS (66–69). And lastly, we assumed that changes in gene expression parallel changes in protein abundance, an assumption that may not always be true (70, 71).

In summary, we have used an integrative approach across scales to examine functional deficits related to metabolism in the brain of a DS mouse model. Our *in vivo* evidence of reduced CBF and CMRO_2_ are concordant with the clinical findings in people with DS, highlighting Dp16 mice as a valuable model to study DS-associated cerebrovascular and metabolic deficits. The physiological changes in the brain of Dp16 mice appear to trigger an adaptive mitochondrial remodeling program across the brain, likely contributing to the preservation of mitochondrial respiratory capacity. Extrapolating from our data, we argue that variable compensations across space and time contribute to variable phenotypic outcomes in DS. Knowledge of the compensatory response may provide pathway-specific targets for therapeutic intervention to improve the quality of life for people with DS.

## MATERIALS AND METHODS

### Mouse model

Dp(16)1Yey/+ (abbreviated Dp16) mice and wild-type (WT) littermate controls, on C57BL/6J genetic background, were obtained from the Jackson Laboratory (Strain # 013530). Mice were fed a standard chow (Envigo; 2018SX). Mice were housed in polycarbonate cages on a 12h:12h light-dark photocycle (lights on at 6 am, lights off at 6 pm) with ad libitum access to water and food. At termination of the study, food was removed 2 h prior to euthanasia. Tissues from different brain regions were collected, snap-frozen in liquid nitrogen, and kept at 80°C until analysis.

All mouse protocols were approved by the Institutional Animal Care and Use Committee of the Johns Hopkins University School of Medicine (animal protocol # MO25M273). All animal experiments were conducted in accordance with the National Institute of Health guidelines and followed the standards established by the Animal Welfare Acts.

### Magnetic resonance imaging (MRI)

All MRI experiments were performed on an 11.7T Bruker Biospec system (Bruker, Ettlingen, Germany) equipped with a horizontal bore and actively shielded pulsed field gradients (maximum intensity of 0.74 T/m) as we have previously described (72–76). All mice were at 16 weeks of age. Images were acquired using a 72-mm quadrature volume resonator as a transmitter and a four element (2 × 2) phased-array coil as a receiver. The B0 field over the mouse brain was homogenized by a global shimming (up to 2nd order) based on a pre-acquired subject-specific field map. The respiration rate was monitored (SA Instruments, Stony Brook, NY, USA) during the experiment to ensure the survival of the mouse. Anesthesia was administered under the following regimen: 1.5% vaporized isoflurane was used for 15 min as the induction followed by a continuous 1.0% isoflurane for maintenance until the end of experiments; during the experiment, the maintenance dosage would be increased slightly to ∼1.2% in cases where a mouse breathed at a rate >150 breaths per minute. At the 10th minute under 1.5% isoflurane inhalation, the mouse was immobilized with a bite bar and ear pins, and placed onto a water-heated animal bed with temperature control before entering the magnet. Each MRI session consisted of MRI measurements detailed below.

### Cerebral oxygen extraction fraction

Cerebral venous oxygenation (Yv) was assessed non-invasively from the confluence of sagittal sinuses with a T2-relaxation-under-spin-tagging (TRUST) technique, which was originally developed on human scanners and recently optimized for animal MRI systems (77). In order to visualize the confluence of sagittal sinuses, an axial time-of-flight (TOF) sequence was first performed with the following parameters: repetition time (TR)/echo time (TE) = 20/2.6 ms, field of view (FOV) = 16 × 16 mm^2^, matrix size = 256 × 256, 35 axial slices, slice thickness = 0.5 mm, and scan duration = 2.2 min. The TRUST scan was then positioned based on the TOF images, and was repeated three times to improve precision. The TRUST sequence used the following parameters: TR/TE = 3500/6.5 ms, FOV = 16 × 16 mm^2^, matrix size = 128 × 128, slice thickness = 0.5 mm, EPI factor = 16, inversion-slab thickness = 2.5 mm, post-labeling delay = 1000 ms, , eTE = 0.13, 10, 20, 40 ms, echo spacing of eTE = 5.0 ms, and scan duration = 3.8 min.. Data processing of TRUST MRI was conducted with a custom written graphic-user-interface (GUI) tool built on MATLAB (MathWorks, Natick, MA) and followed procedures detailed previously (76, 77). Briefly, for each TRUST dataset, subtraction between the control and labeled images was performed to obtain difference images (**Figure 2A**). A region of interest (ROI) was manually drawn on the difference image to encompass the sinus confluence. Four voxels within the ROI with the largest difference signals were automatically selected for spatial averaging. Then, venous blood signal intensities at three different eTEs were fitted into a mono-exponential function to obtain T2. Finally, T2 was converted into Yv using a T2-Yv calibration plot reported by Li et al. (78). Arterial oxygenation is generally close to unity and is assumed to be 0.99 (79). Oxygen extraction fraction (OEF) is defined as the arteriovenous difference in oxygenation and was calculated using *OEF* = 0.99 − *Y*_v_ (80).

### Cerebral Blood Flow

Global cerebral blood flow (CBF) was evaluated with phase-contrast (PC) MRI covering three major feeding arteries, i.e., left internal carotid artery (LICA), right internal carotid artery (RICA), and basilar arteries (BA), in separate scans to collect corresponding through-plane velocity maps (76, 81). Prior to the PC scans, we first performed a coronal TOF angiogram (7 slices, slice thickness = 0.5 mm, no inter-slice gap, TR/TE = 45/2.6 ms, scan duration = 2 min) to visualize the feeding arteries. Then, a sagittal TOF (single slice, tilted to contain the target artery identified from coronal TOF images, thickness = 0.5 mm, TR/TE = 60/2.5 ms, scan duration = 0.4 min) was applied to visualize the in-plane trajectory of the target artery. PC MRI was then positioned using both TOFs and performed using following parameters: TR/TE = 60/3.2 ms, FOV = 15 × 15 mm^2^, matrix size = 300 × 300, slice thickness = 0.5 mm, number of average = 4, dummy scan = 8, receiver bandwidth = 100 kHz, flip angle = 25°, partial Fourier acquisition factor = 0.7, encoding velocity = 20 cm/s, and scan duration = 0.6 min. Processing of the PC dataset was performed with custom-written MATLAB scripts (76, 81). The artery of interest was first manually delineated on the complex-difference image, which shows an excellent contrast between vessel and surrounding tissue. The mask was then applied to the phase velocity map and the integration of arterial voxels yields blood flow through that targeted artery in ml/min. Summation of blood-flow values across the four major feeding arteries yielded total blood flow to the brain. To further account for the brain-size differences and to obtain unit-mass CBF values, the total blood flow was divided by the brain weight, which was calculated as the product of brain volume and density [1.04 g/ml (82)]. A fast spin-echo MRI sequence was used to assess brain volume under the following parameters: TR/TE = 4000/10.0 ms, field of view = 15× 15 mm^2^, matrix size = 128 × 128, slice thickness = 0.5 mm (without inter-slice gap), echo spacing = 5.0 ms (4 spin echoes per scan), 35 axial slices, and scan time = 2.1 min (83). The global CBF value is written in the unit of milliliters per 100 g brain tissue per minute (ml/100g/min). Inter-rater reproducibility of PC MRI processing has previously been assessed and reported an interrater correlation of >95% (81).

### Cerebral metabolic rate of oxygen

Cerebral metabolic rate of oxygen (CMRO_2_) was computed from Yv and CBF using the Fick principle, i.e., CMRO_2_ = Ca · OEF · CBF, where Ca denotes the molar concentration of oxygen in a unit volume of blood and was assumed to be 882.1 µmol O2/100 ml blood based on previous literature (84). CMRO_2_ is written in the unit of µmol oxygen per 100 g brain tissue per min (µmol O_2_/100 g/min).

### Mitochondrial respirometry

Respirometry was conducted on frozen tissue samples to assay for mitochondrial activity as we have previously described (46, 47). Briefly, the cerebellum, cortex, hypothalamus and hippocampus were dissected from mice 21 weeks of age, snapped frozen in liquid nitrogen, and stored at -80°C for later analysis. Samples were thawed in MAS buffer (70mM sucrose, 220 mM mannitol, 5 mM KH2PO4, 5 mM MgCl2, 1 mM EGTA, 2 mM HEPES pH 7.4), finely minced with scissors, and then homogenized with a glass dounce homogenizer. The resulting homogenate was spun at 1000 g for 10 min at 4°C. The supernatant was collected and immediately used for protein quantification by BCA assay (Thermo Scientific, 23225). Each well of the Seahorse microplate was loaded with 6 µg of cortex, cerebellum, hippocampus and hypothalamus. Each biological replicate consists of three technical replicates. Samples from all tissues were treated separately with NADH (1 mM) as a complex I substrate or Succinate (a complex II substrate, 5 mM) in the presence of rotenone (a complex I inhibitor, 2 µM), then with the inhibitors rotenone (2 µM) and Antimycin A (4 µM), followed by TMPD (0.45 mM) and Ascorbate (1 mM) to activate complex IV, and finally treated with Azide (40 mM) to assess non-mitochondrial respiration. Additionally, to isolate CV-dependent activity, three baseline time points were first recorded. All samples were then treated with 4 μM rotenone (Complex I inhibitor) and 4 μM antimycin A (Complex III inhibitor), followed by three additional measurement time points. Complex V was subsequently activated by the addition of 20 mM ATP (Sigma A26209) together with 7.87 mM carbonyl cyanide p-trifluoromethoxyphenylhydrazone (FCCP; Enzo BML-CM120), an uncoupler of oxidative phosphorylation, and three more time points were collected. Finally, 10 μM oligomycin (Sigma 495455) was added to inhibit Complex V and distinguish non-mitochondrial acidification. All mitochondrial respiration data were normalized to mitochondrial content, quantified using MitoTracker Deep Red (MTDR, ThermoFisher, M22426) as described (insert new numbers). Briefly, lysates were incubated with MTDR (1 µM) for 10 min at 37°C, then centrifuged at 2000 g for 5 min at 4°C. The supernatant was carefully removed and replaced with 1x MAS solution and fluorescence was read with excitation and emission wavelengths of 625 and 670 nm, respectively. To minimize non-specific background signal contribution, control wells were loaded with MTDR and 1x MAS and subtracted from all sample values.

### RNA-sequencing and bioinformatics analysis

Bulk RNA sequencing of mouse cortex, hippocampus, cerebellum, and hypothalamus were performed by Novogene (Sacramento, California, USA) on a NovaSeq X Plus platform and pair-end reads (2 x 150 bp) were generated, with 6 G raw data per sample. Mice were 21 weeks of age when brain tissues were collected. Sample size for cortex, hippocampus, and cerebellum: *n* = 8 per sex per genotype per tissue. Sample size for hypothalamus: *n* = 6 per sex per genotype per tissue. A total of 72 tissue samples were subjected to RNA sequencing. We have previously published the RNA sequencing data for the hypothalamus (21) but not the other brain regions. All sequencing data were analyzed using the standard Novogene Analysis Pipeline. Sequencing reads were aligned to Mus musculus reference genome (GRCm39/mm39). Data analysis was performed using a combination of programs, including Fastp, Hisat2, and FeatureCounts. Differential expressions were determined through DESeq2. The resulting *P*-values were adjusted using Benjamini and Hochberg’s approach for controlling the false discovery rate (FDR). Genes with an adjusted *P*-value (Padj) ≤ 0.05 found by DESeq2 were assigned as differentially expressed. All DEGs up- and down-regulated in cortex, hippocampus, cerebellum, and hypothalamus are provided in **supplemental Table 1-8**. Gene ontology (GO), Kyoto Encyclopedia of Genes and Genomes (KEGG), and Reactome (http://www.reactome.org) enrichment were implemented by ClusterProfiler. All volcano plots and heat maps were generated in Graphpad Prism 10 software. All statistics were performed on log transformed data. All heat maps were generated from column z-score transformed data. The z-score of each column was determined by taking the column average, subtracting each sample’s individual expression value by said average then dividing that difference by the column standard deviation: Z-score = (value – column average)/column standard deviation. High-throughput sequencing data from this study have been submitted to the NCBI Sequence Read Archive (SRA) under accession number # PRJNA1160420.

### Modular gene expression analysis across brain regions and sex

Mitochondrial gene expression modules were defined using mitochondrial localization and pathway-level classifications from the Mouse MitoCarta3.0 database (85). All gene symbols are provided in **Supplemental table 9**. Normalized expression values for each gene were Z-score transformed on a gene-wise basis across individual mice within each sex and brain region. Module scores were then calculated for each individual mouse by averaging the normalized expression values of all genes assigned to the corresponding module. Thus, each mouse was represented by a module-level expression score for each mitochondrial pathway.

For each brain region, mitochondrial module scores were compared between WT and Dp16 mice within sex. The Δmodule score was calculated as the difference in mean module score between Dp16 and WT mice, with positive values indicating greater module expression in Dp16 mice and negative values indicating greater module expression in WT mice. Individual mouse module scores were retained for visualization and statistical analysis. Statistical significance was assessed using unpaired Welch’s *t*-tests, followed by Holm-Sidak correction for multiple comparisons. Adjusted significance values were utilized for analysis, with significance represented graphically by circle size as shown in **Figure 8A**.

An integrated multi-region mitochondrial module analysis was performed to provide a composite representation of mitochondrial transcriptional remodeling across the brain. For each sex and genotype, the corresponding module score was averaged across all analyzed brain regions, with each brain region weighted equally, to generate an integrated multi-region module score. The integrated Δmodule score was calculated as the difference between the Dp16 and WT integrated module scores. Positive values indicate greater integrated module expression in Dp16 mice, whereas negative values indicate greater expression in WT mice. Because this analysis was intended to visualize broad trends in mitochondrial remodeling across regions rather than provide an independent statistical test, no significance testing was performed on the integrated values for **Figure 8B**.

### Secondary gene expression module analysis

Secondary gene expression modules were developed to contextualize the mitochondrial remodeling identified in **Figure 8A-B** by examining biological processes that may contribute to, accompany, or provide additional context for altered mitochondrial function. Secondary modules included cytoplasmic translation, mitochondrial protein import, mitochondrial proteostasis, mitochondrial translation, mitochondrial tRNA charging, mitophagy, redox homeostasis, neuronal activity, neurovascular coupling, vascular development/angiogenesis, and vascular function/endothelial signaling. Gene symbols and module classifications for each secondary module are provided in **Supplemental Table 10**.

Secondary module scores were calculated using the same gene-wise Z-score normalization and module-scoring approach described above. For each mouse, normalized expression values for genes assigned to a given secondary module were averaged to generate a module-level expression score. Module scores were compared between Dp16 and WT mice within each sex and brain region. The Δmodule score represents the difference between the mean module score in Dp16 and WT mice, with positive values indicating greater expression in Dp16 mice and negative values indicating greater expression in WT mice.

To summarize the relative magnitude of secondary module remodeling across brain regions, the absolute value of each Δmodule score was calculated to quantify the magnitude of remodeling independent of direction. Absolute values were used only to quantify and rank the magnitude of module-level remodeling independent of direction; the underlying module scores and Δmodule scores were calculated without taking absolute values, such that the direction of Dp16 versus WT change was retained. For each brain region, the cumulative magnitude of remodeling across secondary modules was calculated as the sum of absolute Δmodule scores (Σ|**Δ**module score|). Brain regions were ranked according to this cumulative value. Within each brain region, individual secondary modules were ranked according to their relative absolute Δmodule score, such that modules with larger absolute differences were represented by larger graphical elements as shown in **Figure 8C**. The signed Δmodule score was retained for interpretation of direction, with positive Δmodule scores indicating greater expression in Dp16 mice and negative values indicating greater expression in WT mice.

Statistical significance of secondary module differences was assessed using unpaired Welch’s *t*-tests followed by Holm-Sidak correction for multiple comparisons. Statistical significance was represented in **Figure 8C** by the width of the corresponding feature in the “Magnitude and Significance” column. Significance categories included non-significant (ns) and nominally suggestive differences (uncorrected *P*<0.05).

### Genome-wide differential expression and shared transcriptional program analysis

To identify transcriptional features shared between brain regions exhibiting prominent mitochondrial remodeling, genome-wide differential expression was examined independently of the predefined mitochondrial and secondary gene expression modules. Differential expression was assessed between Dp16 and WT mice within the male cortex and female cerebellum, representing the regions with the greatest mitochondrial module remodeling in the respective sexes. Genes were considered differentially expressed at |Log_2_FC|>1.0 and the false discovery rate (FDR)-adjusted *P* value was <0.05.

DEGs identified independently within the male cortex and female cerebellum were separated according to direction of change, defined as up-regulated and down-regulated in Dp16 relative to WT. Genes exhibiting concordant differential expression in both regions were identified by intersection of the corresponding direction-specific DEG lists. This yielded 642 genes commonly up-regulated and 313 commonly down-regulated in Dp16 relative to WT across male cortex and female cerebellum. Genes identified in the overlap analysis, including their directional classifications, are provided in **Supplemental Table 11**.

To determine the biological processes represented by the shared transcriptional programs, the concordantly up-regulated and down-regulated gene sets were analyzed independently using Enrichr (86). Functional enrichment was performed separately for the shared-up and shared-down gene sets to preserve the direction of transcriptional change. The Gene Ontology Biological Process 2026 (GOBP), Cellular Component 2026 (GOCC), and Molecular Function 2026 (GOMF) libraries were evaluated. Enrichment significance was assessed using multiple-testing-adjusted *P* values reported by Enrichr. A total of 44 significantly enriched terms were identified among the shared-up genes after multiple-testing correction, whereas no terms reached statistical significance among the shared-down genes. All significantly enriched terms are provided in **Supplemental Table 12**.

### Statistical analyses

All results are expressed as mean ± standard error of the mean (SEM). Statistical analysis was performed with GraphPad Prism 10 software (GraphPad Software, San Diego, CA). Data were analyzed with two-tailed Student’s *t*-tests or by repeated measures ANOVA. For two-way ANOVA, we performed Sidek or Bonferroni post hoc tests. *P* < 0.05 was considered statistically significant.

## AUTHOR CONTRIBUTION

GWW and MMS: Conceptualization; MS, DCS, ZW, GWW: Formal analysis; GWW, ZW: Funding acquisition; MS, DCS, ZW, GWW: Investigation; ZW: Methodology; GWW: Project administration; GWW: Supervision; MMS, DCS, ZW, GWW: Visualization; GWW, MMS, DCS: Roles/Writing - original draft; DCS, ZW: Writing - review & editing.

## ACKNOWLEDGEMENTS

The work was funded, in part, by grants from the National Institute of Health (DK084171 to GWW; AG081932 to ZW). The MRI equipment in this study was funded by National Institute of Health grant S10 OD032188

## DATA AVAILABILITY

All RNA-seq data have been deposited in NCBI Sequence Read Archive (SRA), under BioProject # PRJNA1463876. All raw scan data from MRI are available upon request.

## SUPPLEMENTAL FIGURE FILES LEGENDS AND SOURCE DATA

**Supplemental figure 1.** Mitochondrial content as judged by MTDR levels across brain regions in males (A) and females (B) mice. Sample size per sex: WT = 8 and Dp16 = 8.

**Supplemental figure 2.** Mitochondrial modules across brain regions in male and female mice. All of the mitochondrial module values per mouse across sex and genotype used for figure 8A-B.

**Supplemental figure 3.** Secondary modules across brain regions for male mice. All secondary module values per male mouse across brain regions and genotypes.

**Supplemental figure 4.** Secondary modules across brain regions for female mice. All secondary module values per female mouse across brain regions and genotypes.

**Supplemental Table 1**. All differential expressed genes (DEGs) in the cortex of Dp16 male mice.

**Supplemental Table 2**. All differential expressed genes (DEGs) in the cortex of Dp16 female mice.

**Supplemental Table 3**. All differential expressed genes (DEGs) in the hippocampus of Dp16 male mice.

**Supplemental Table 4**. All differential expressed genes (DEGs) in the hippocampus of Dp16 female mice.

**Supplemental Table 5**. All differential expressed genes (DEGs) in the cerebellum of Dp16 male mice.

**Supplemental Table 6**. All differential expressed genes (DEGs) in the cerebellum of Dp16 female mice.

**Supplemental Table 7**. All differential expressed genes (DEGs) in the hypothalamus of Dp16 male mice.

**Supplemental Table 8**. All differential expressed genes (DEGs) in the hypothalamus of Dp16 female mice.

**Supplemental Table 9.** All gene symbols for mitochondrial module classification list used in figure 8A. All genes and their mitochondrial module classification used for figure 8A.

**Supplemental Table 10.** All gene symbols for secondary module classification list used in figure 8C. All genes and their secondary module classification used for figure 8C.

**Supplemental Table 11.** All gene symbols for DEG overlap classification used in figure 8D. All directionally matched overlapping genes between male cortex and female cerebellum used for figure 8D.

**Supplemental Table 12.** Male cortex and female cerebellum shared DEG pathway enrichment for figure 8D. All enriched pathways of male cortex and female cerebellum shared differentially expressed genes used for figure 8D.

